# A small molecule inhibitor Mirin prevents TOP3A-dependent mtDNA breakage and segregation

**DOI:** 10.1101/2024.03.14.585071

**Authors:** Koit Aasumets, Anu Hangas, Cyrielle P. J. Bader, Direnis Erdinc, Sjoerd Wanrooij, Paulina H. Wanrooij, Steffi Goffart, Jaakko L.O. Pohjoismäki

**Affiliations:** Department of Environmental and Biological Sciences, University of Eastern Finland, P.O. Box 111, 80101 Joensuu, Finland; Department of Medical Biochemistry and Biophysics, Umeå University, 901 87 Umeå, Sweden; Department of Medical Biochemistry and Cell Biology, University of Gothenburg, Gothenburg SE-40530, Sweden

**Keywords:** MGME1, mitochondrial DNA, DNA topology, DNA replication, double-strand break, DAMP, inflammation, molecular biology

## Abstract

Mirin, the chemical inhibitor of MRE11, has been recently reported to prevent immune response activation caused by mitochondrial DNA (mtDNA) breakage and release upon replication stalling. We show here that Mirin prevents mitochondrial replication fork breakage in mitochondrial 3’-exonuclease MGME1 deficient cells and the resulting innate immune response induction, but that this occurs independently of MRE11. Furthermore, Mirin also caused alteration of mtDNA supercoiling and accumulation of hemicatenated replication termination intermediates, hallmarks of topoisomerase dysfunction, as well as alleviated topological changes induced by the overexpression of mitochondrial TOP3A, including TOP3A-dependent strand breakage at the non-coding region of mtDNA, potentially explaining its protective effect in the MGME1-knockout cells. Although Mirin does not inhibit TOP3A *in vitro*, our results demonstrate its MRE11-independent effects in cells and give insight into the mechanisms of mtDNA segregation, as well as the maintenance of genomic integrity in mitochondria.

**Significance Statement:**

- Broken mitochondrial DNA (mtDNA) in *MGME1* knockout cells activates innate immune response, which is prevented by Mirin, a small molecule inhibitor of MRE11.
- Mirin also interferes with mtDNA replication termination and segregation, suggesting that termination intermediates or paused forks are a major source of mtDNA breakage.
- We show that these effects are likely dependent on topoisomerase 3A (TOP3A) -related processes in mitochondria, questioning the Mirin target also in the nucleus.

## Introduction

Mitochondria are essential cell organelles producing the majority of cellular ATP through oxidative phosphorylation, as well as being involved in various biosynthetic processes, cell signaling, apoptosis and more (McBride *et al*., 2006). As a relic of their past as free-living proteobacteria-like organisms (Martijn *et al*., 2018), mitochondria have their own genome, a circular double-stranded DNA ring. Mitochondrial DNA exists in cells in hundreds to thousands of copies and can take a number of different topological conformations, including covalently closed circles, open circles, supercoils, monomeric dimers as well as various catenated forms (Goffart *et al*., 2019). MtDNA topology is controlled by topoisomerases, of which TOP3A is required for the release of torsional stress during replication (Hangas *et al*., 2022) as well as resolution of hemicatenane intermediates (Nicholls *et al*., 2018) that form on the non-coding region (NCR) of the genome upon replication termination.

Prior to its recently discovered mitochondrial functions, the nuclear roles of TOP3A have been studied excessively. Besides removing hemicatenanes arising from converging nuclear DNA replication forks (Lee *et al*., 2019), TOP3A is involved in resolution of recombination junctions (Wu *et al*., 2006), assisting replication restart (Shorrocks *et al*., 2021) as well as introducing positive supercoiling to facilitate sister chromatid separation in mitosis (Bizard *et al*., 2019). Unlike in mitochondria, where TOP3A appears to function alone, its nuclear roles require partnering with the OB-fold domain proteins RMI1 and RMI2, as well as a helicase, such as BLM (BLM-TOP3A-RMI1-RMI2 or BTR complex) in the case of recombination resolution (Xu *et al*., 2008), FANCM and BTR at stalled forks (Deans and West, 2009) as well as PICH during mitosis (Bizard *et al*., 2019).

Another complex involved in rescue of stalled forks in the nucleus is the MRE11-RAD50-NBS1. Besides regulating double-strand break (DSB) repair (Bressan *et al*., 1999), the MRE11 complex stabilizes components of the replisome (Tittel-Elmer *et al*., 2009) and promotes replication restart through homology-dependent repair (HDR) (Trenz *et al*., 2006). Lack of MRE11 results in collapse of the stalled fork and generation of DSBs. Similarly to the nucleus, replication stress in mitochondria can also result in double-strand breaks, generating linear mtDNA fragments, which often consist of the region between the two replication origins, OriH and OriL (Torregrosa-Munumer *et al*., 2019). In fact, mitochondria have an efficient enzymatic machinery to turn over broken mtDNA, relying on the 5’-exonuclease MGME1, the 3’-exonuclease activity of mitochondrial replicative polymerase Pol γ and TWNK helicase (Peeva *et al*., 2018).

Unless degraded, fragmented mtDNA has the tendency to exit mitochondria and enter the cytoplasm, where it can act as a damage-associated molecular pattern (DAMP) to elicit inflammation either *via* the endosomal Toll-like receptor–dependent response (Bao *et al*., 2016), the inflammasome (Zhong *et al*., 2018), cGAS/STING-pathway (West *et al*., 2015) or RIG-I (Tigano *et al*., 2021). Recently, it was shown that Mirin, a chemical inhibitor of MRE11 (Dupre *et al*., 2008; Garner *et al*., 2009), was able to block the immune response caused by mtDNA replication stress, indicating a role for MRE11 in preventing replication fork breakage in mitochondria (Luzwick *et al*., 2021). Inspired by this finding, we sought to see whether MRE11 could influence the abundance of the OriH-OriL linear mtDNA fragments present in *MGME1* knockout (KO) HEK293T cells. We previously suggested that these fragments arise naturally from frequent breakage at these replication origins due to prolonged pausing and that their accumulation in *MGME1* KO cells is due to the defective linear DNA turnover machinery and not due to a role of MGME1 in replication *per se* (Torregrosa-Munumer *et al*., 2019).

While Mirin treatment effectively decreased the levels of linear mtDNA and abolished the immune response in the *MGME1* KO cells, we were unable to confirm the effect to be dependent on MRE11 inhibition. Instead, Mirin induced topological changes and accumulation of hemicatenated mtDNA, which are consistent with the inhibition of TOP3A functions in mitochondria. Furthermore, while Mirin did not inhibit TOP3A activity *in vitro*, it was able to counteract the effects of mitochondrially targeted TOP3A overexpression, including relaxation of supercoiled mtDNA and its breakage within the NCR. The results suggest that TOP3A-dependent fork breakage in the NCR explains the accumulation of broken mtDNA in MGME1-knockout cells, as well as give insight into the resolution of mitochondrial replication termination intermediates.

## Results

As mitochondrial double-strand breaks have been reported to elicit immune reaction in cultured cells (Tigano *et al*., 2021), we wanted to see if *MGME1* KO cells, possessing abundant linear mtDNA fragments, showed activation of the inflammatory cytokine response as indicated by STAT1 phosphorylation (Pilz *et al*., 2003). As expected, *MGME1* KO cells showed chronic STAT1 activation (Figure 1A), confirming that the broken mtDNA, in this case mostly consisting of aborted replication intermediates spanning between OriH and OriL (Peeva *et al*., 2018; Torregrosa-Munumer *et al*., 2019), can escape the mitochondrial compartment to function as a DAMP. This STAT1 activation was almost completely abolished by Mirin treatment, confirming the previous findings (Luzwick *et al*., 2021) and correlated with significantly reduced levels of the OriH-OriL fragment (Figure 1B, C). However, the knockdown of MRE11 did not phenocopy the Mirin effects on STAT1 activation nor the levels of linear mtDNA (Figure S1A, B). In fact, we were unable to find evidence of the mitochondrial localization of MRE11 in *MGME1* KO cells using protease protection assay (Figure S1C) or confocal microscopy (Figure S1D). Similarly, a recent study on *Drosophila* found no MRE11 from mitochondria (Klucnika *et al*., 2023).

**Figure 1.**
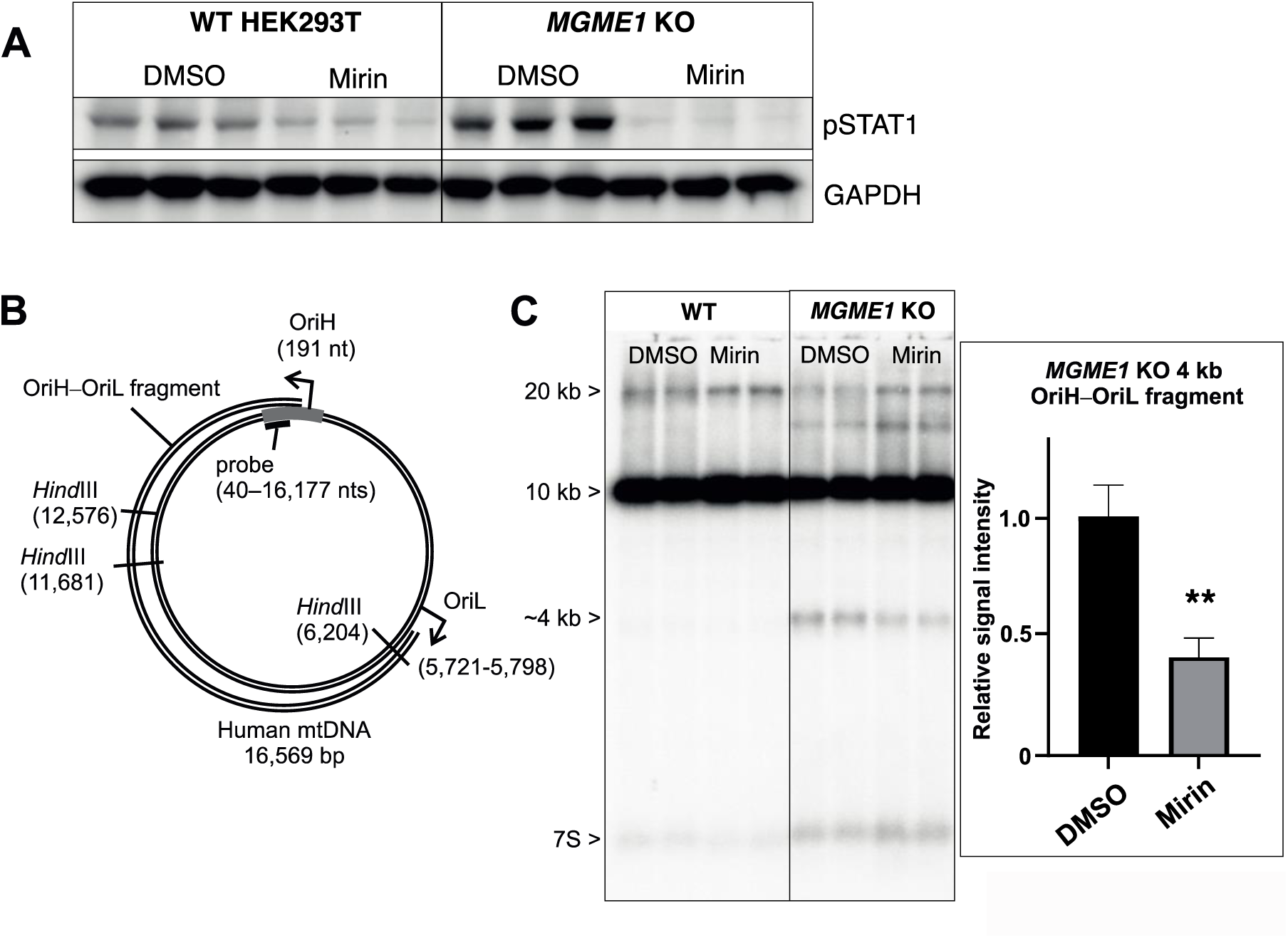
Mirin inhibits STAT1 activation and mtDNA breakage in *MGME1* knockout cells. (A) *MGME1* knockout (KO) cells show immune response activation, indicated by the elevated levels of phosphorylated STAT1. This activation is almost completely abolished by a 48 h treatment with 100 μM Mirin. (B) Schematic illustration of human mtDNA, showing the OriH-OriL fragment present in *MGME1* KO cells as well as the location of *Hind*III restriction sites and the NCR probe used for Southern blot hybridization. (C) Southern blot of *Hind*III-digested DNA from wildtype (WT) and *MGME1* KO cells, probed with the NCR probe. Note the increase in the dimeric 20 kb DNA species as well as the significant reduction in the 4 kb OriH-OriL fragment abundance after 48 h of 100 μM Mirin. Ratios of 4 kb OriH-OriL /10 kb fragment from 4 independent experiments are shown with standard error of the mean. Statistical significance (*P-*value 0.0015) was calculated with two-tailed Student’s t-test.

Besides the decrease in OriH-OriL fragment in the *MGME1* KO cells, the appearance of an apparent dimeric (2n) mtDNA species (20 kb in Figure 1C) in all Mirin-treated samples caught our attention. Two-dimensional agarose gel electrophoresis (2D-AGE) investigation of the NCR-containing parts of the mtDNA revealed that this band represented termination intermediates (Wanrooij *et al*., 2007; Nicholls *et al*., 2018), whose abundance was increased in Mirin-treated cells together with fully double-stranded replication forks, especially at the descending part of the y-arc (Figure 2). While the accumulation of replication forks indicates impaired progression of replication, the termination intermediates represent a very specific molecular class, namely hemicatenanes, which can only be resolved by TOP3A in mitochondria (Nicholls *et al*., 2018). Interestingly, also the increase in the descending part of the y-arc is similar to the one seen in TOP3A knockdown (Nicholls *et al*., 2018).

**Figure 2.**
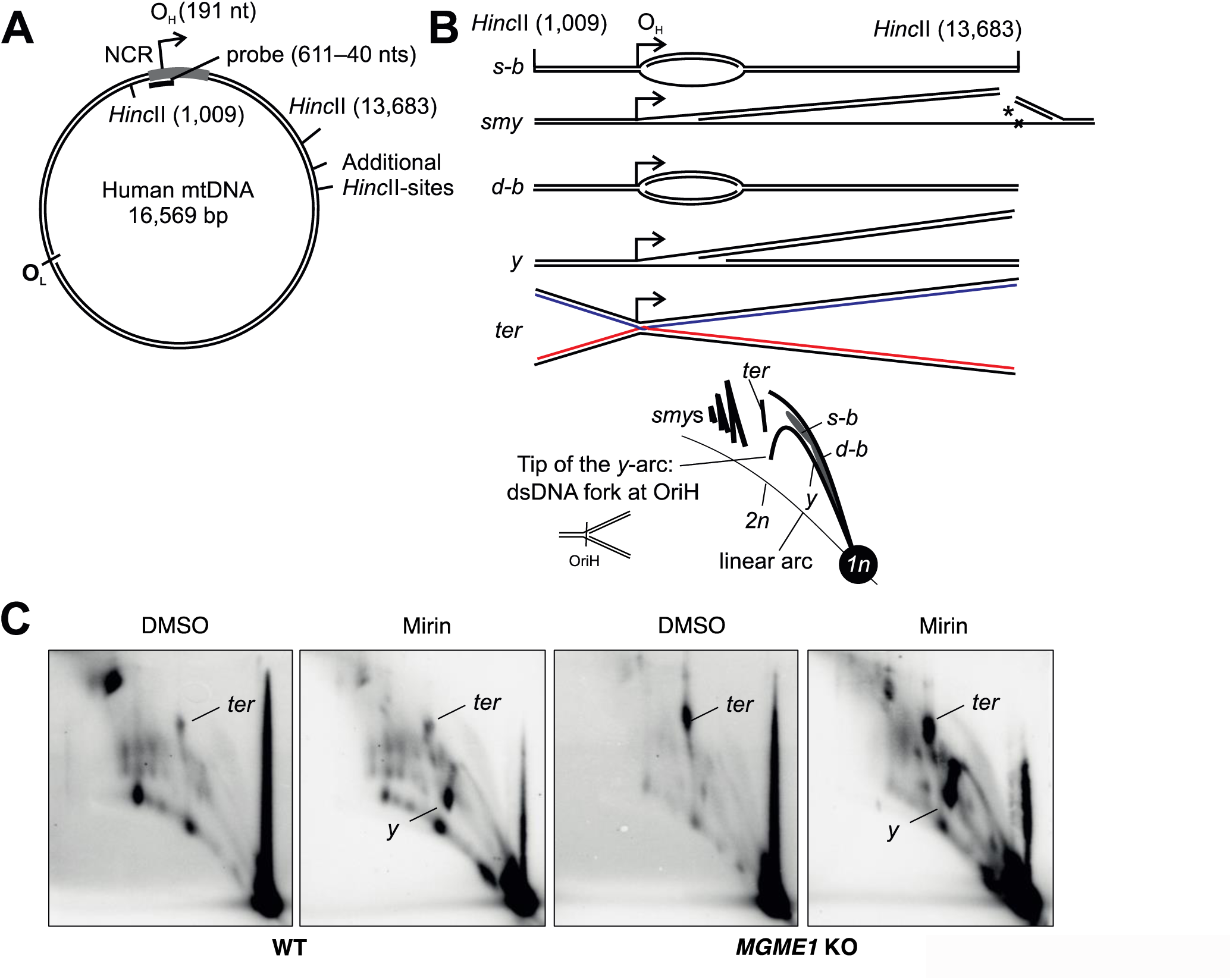
Increased levels of mtDNA replication termination intermediates after Mirin treatment. (A) Schematic illustration of human mtDNA showing restriction cut sites and the probe locations. Details of non-relevant cut sites are omitted for clarity. (B) Replication intermediates present in the OriH-containing *Hinc*II fragment and their migration patterns on 2D-AGE. Single-stranded bubble arc (*s-b*) represents partially single-stranded (ss)DNA replication bubbles. Slow moving y-forms (*smy*) are partially ssDNA replication forks that have progressed beyond the downstream restriction sites (*), giving rise to high-molecular weight replication intermediates. Double-stranded (ds)DNA replication intermediates are represented by proper bubble (*d-b*) and y-arcs (*y*). Note that the y-arc is not complete because of the position of OriH within the fragment; its descending tip corresponds to persistent replication forks at this locus. Four-way junctional termination intermediates (*ter*) form at the end of replication close to the OriH. (C) 48 h 100 μM Mirin treatment of cells increases the termination intermediates (*ter*) as well as replication forks paused at OriH (*y*).

To see if the Mirin-treated cells had other phenotypes that might be related to topoisomerase dysfunction, we next looked at mtDNA topology. To our surprise, Mirin treatment resulted in an increase in the well-bound mtDNA signal, which represents complex DNA unable to migrate into the gel, concomitant with an apparent reduction in the more clearly defined catenane bands (Figure 3A). Besides open monomeric circles (OC), mtDNA in HEK293 cells typically also consists of supercoiled species, which run as a defined band (SC) due to its constant linking number (Hangas *et al*., 2018). Interestingly, the wildtype HEK293T cells treated with Mirin showed additional, more heterogeneous supercoiled forms (SC* in Figure 3A), which were absent in *MGME1* KO cells. We also noted a notable decrease in TOP3A protein levels after 48h Mirin treatment (Figure 3B).

**Figure 3.**
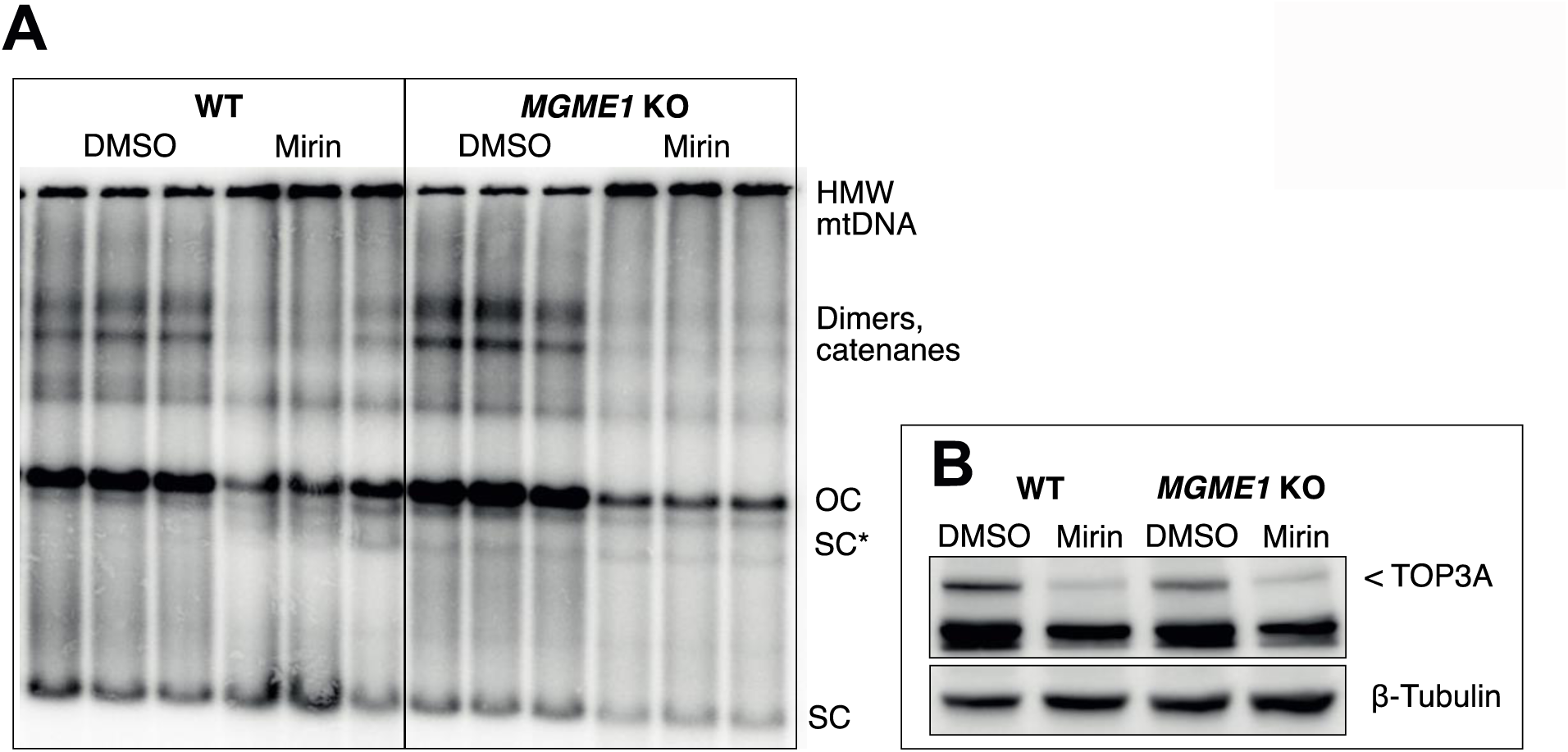
Mirin treatment alters mtDNA topology. (A) A 48 h 100 μM Mirin treatment of wildtype (WT) HEK293 cells causes increase in fully supercoiled (SC) mtDNA and generates a ladder of additional supercoiled forms (SC*) below the fully relaxed open circles (OC). The effect on supercoiling is not as dramatic in *MGME1* KO cells. (B) Mirin treatment reduces TOP3A protein levels in the same cells. Unspecific band under the TOP3A band is shown together with β -tubulin loading control. See also the Figure S4C for the confirmation of the antibody specificity.

As we have previously shown that overexpression of mitochondrially targeted TOP3A (mtTOP3A) results in excess mtDNA breakage due to strand cleavage at or close to the OriH as well as loss of supercoiled forms (Hangas *et al*., 2022), we were curious to see whether Mirin treatment could compensate these effects. Indeed, 48 h Mirin treatment was able to partially rescue the mtTOP3A expression induced resolution of the HMW forms as well as the loss of supercoiled mtDNA (Figure 4A, B). Furthermore, Mirin reduced the mtTOP3A-dependent cleavage of termination intermediates (Figure 4C, D, Figure S2). The association of termination intermediate accumulation with mtDNA segregation defect was corroborated by the observed aggregation of mtDNA-nucleoids in the Mirin treated cells (Figure S3).

**Figure 4.**
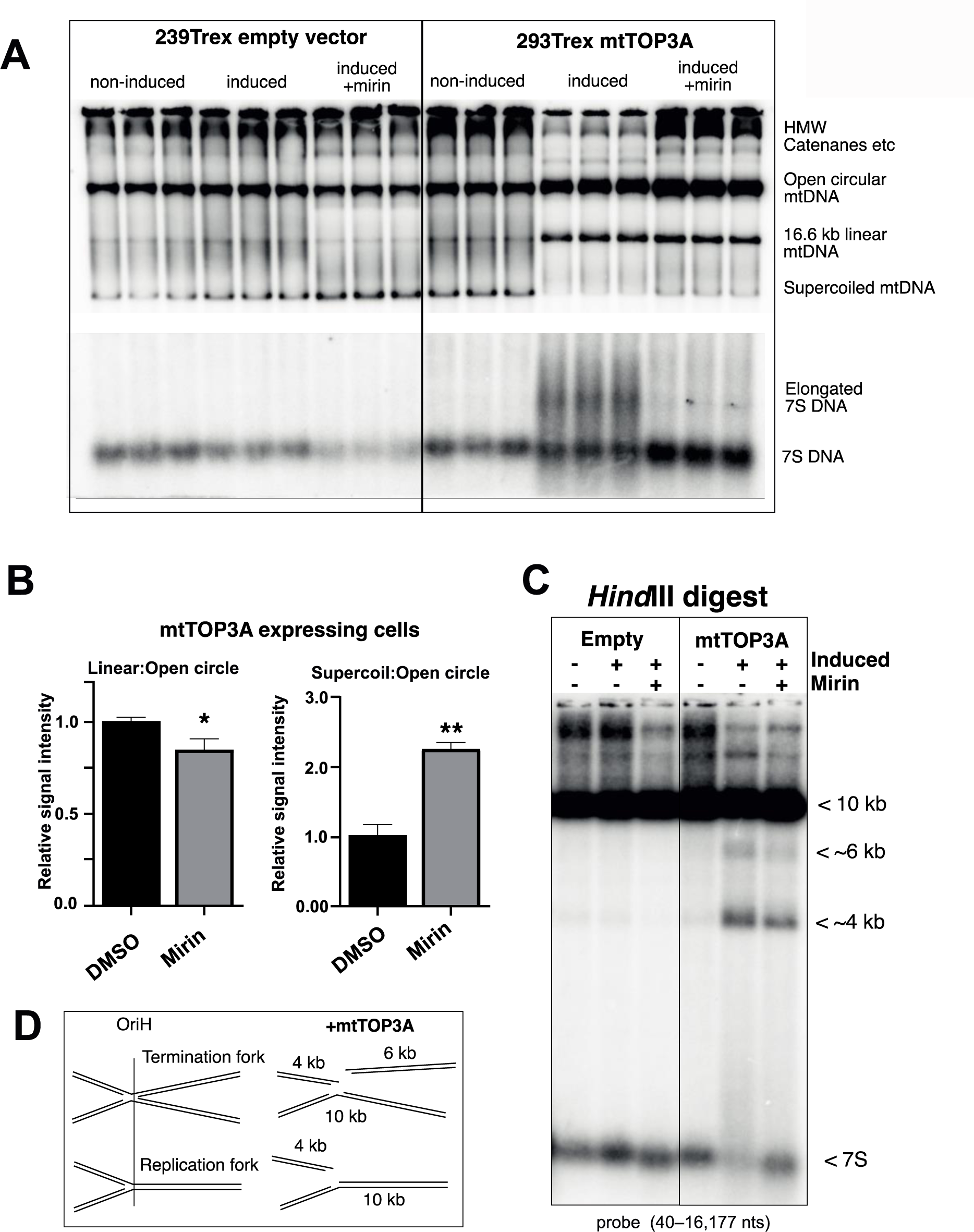
Mirin treatment partially reverses the effects of mitochondrially targeted TOP3A (mtTOP3A) overexpression. (A) A mtDNA topology gel from 293Trex cell lines with an empty vector insertion or tetracyclin inducible mtTOP3A construct showing the effects of transgene induction with and without 100 μM Mirin for 48 h. The bottom panel shows the 7S signal of the same gel. Note the accumulation of linear mtDNA after mtTOP3A induction, the increase in supercoiled mtDNA after Mirin treatment (compare to Figure 3) as well as the lengthening of the 7S DNA. (B) Quantitation of the abundance of linear and supercoiled mtDNA in mtTOP3A expressing cells after 48 h of 100 μM Mirin. (C) Mirin treatment decreases the mtTOP3A-induced breakage in the NCR. Compare with Figure 1C. (D) Schematic illustration of the *Hind*III NCR fragment analyzed in (C). The difference in intensity of the 4 kb and 6 kb fragments is likely due to the probe overlapping the latter only partially.

We have previously shown that mtTOP3A expression results in elongation of the 7S DNA (Hangas *et al*., 2022), a replication intermediate that has terminated soon after initiation and forms the so-called D-loop structure of mtDNA (Jemt *et al*., 2015). Interestingly, Mirin abolished this elongation of the 7S DNA, while increasing its overall abundance in mtTOP3A expressing cells (Figure 4A, bottom panel). The effect of Mirin on 7S DNA is very similar to the knockdown of TOP3A (Figure S4A), although interestingly TOP3A knockdown increased the abundance of OriH-OriL fragments in *MGME1* KO cells in contrast to their decrease under Mirin treatment (Figure S4B *vs.* Figure 1B, C). If the accumulation of the OriH-OriL fragments is associated with replication stalling, as demonstrated before (Torregrosa-Munumer *et al*., 2019), the result would suggest that the fork breakage activity of TOP3A is independent of the removal of torsional stress during replication. In fact, Mirin did not show any inhibitory effect on the TOP3A ability to remove supercoils *in vitro* (Figure S5).

DNA-intercalating drugs can cause alterations in mtDNA topology (Young, 2017; Hangas *et al*., 2018; Hangas *et al*., 2022); and although such a broad spectrum of observed mtDNA effects would be unprecedented as a drug artefact, we tested whether Mirin alone could influence the electrophoretic properties of circular DNA *in vitro*. In contrast to known DNA-intercalating drugs, such as Doxorubicin (Hangas *et al*., 2018) and the telomerase inhibitor RHSP4 (Cookson *et al*., 2005), which altered plasmid migration in ethidium bromide-free gel electrophoresis already at concentrations of 3.4 µM and 0.5 µM, respectively, 100 µM Mirin had no effect (Figure S6).

## Discussion

The ability of the broken mtDNA to escape mitochondria and function as DAMPs is intriguing and likely plays a physiological role in signaling cellular damage (West *et al*., 2015; Tigano *et al*., 2021). As reported here for the first time, the persistent broken replication intermediates in *MGME1 KO* cells seem to have the same function as DAMPs as other types of linear mtDNA fragments (Bao *et al*., 2016; Zhong *et al*., 2018; Luzwick *et al*., 2021; Tigano *et al*., 2021), although in a chronic manner, making these cells a promising model system to study mtDNA release and recognition in the future. Furthermore, we confirmed that Mirin, a small molecule inhibitor of the MRE11 nuclease and suggested to promote replication fork protection in mitochondria (Luzwick *et al*., 2021), can prevent this immune response (Figure 1A). This effect was concomitant with an increase in mtDNA replication termination intermediates as well as accumulation of replication intermediates paused in the NCR (Figure 2). The accumulation of replication intermediates was especially pronounced in the *MGME1* KO cells. As this effect was coinciding with the decreased levels of OriH-OriL fragments in the same cells (Figure 1), the most plausible explanation is that Mirin prevents the resolution of these replication intermediates, which would be in agreement with the proposed role of MRE11 in mitochondria (Luzwick *et al*., 2021).

However, these effects together with the influence of Mirin on mtDNA topology (Figures 2– 4) were not consistent with MRE11 knockdown (Figure S1), which produced no effects on mtDNA whatsoever. The lack of effect is difficult to explain through potential redundancy in the replication fork resolution mechanisms unless Mirin has additional targets in mitochondria. Our observations led us to investigate TOP3A, as it has a vital role in the resolution of the termination intermediates (Nicholls *et al*., 2018; Hangas *et al*., 2022), and whose inhibition could explain the accumulation of termination intermediates (Figure 2C) as well as aggregation of mitochondrial nucleoids (Figure S3), indicative of segregation defect, resulting from Mirin treatment. Mirin also induced general changes in the mtDNA topology, such as abnormal supercoiling (Figure 3). In the light of these observations, it is plausible that interference of TOP3A-related functions might also explain the effect of Mirin on nuclear DNA-related pathways. For example, although MRE11 initiates resection (Shibata *et al*., 2014), which is required for the licensing of homologous recombination (HR) in the nucleus, the TOP3A containing BTRR complex is required for long-range resection during the initial stages of HR (Soniat *et al*., 2023) and only later has a pivotal role in branch migration and resolution of Holliday junctions (Wu *et al*., 2006). Inhibition of either of the initial stages of HR might be difficult to distinguish mechanistically and would nevertheless be consistent with Mirin abolishing the G2/M checkpoint and homology-dependent repair in mammalian cells, as reported initially (Dupre *et al*., 2008). Curiously, none of the known nuclear partners of TOP3A were found in a BioID assay using a mitochondrially-targeted recombinant TOP3A as a bait (Hangas *et al*., 2022), although for example the members of the mitochondrial replisome were identified.

The ability of Mirin to counteract the mtDNA breakage caused by mitochondrially targeted TOP3A (Figure 4) provides interesting insight into the formation of the OriH-OriL fragment in the *MGME1* KO cells and mtDNA replication in general. As explained, besides resolving the hemicatenanes formed during replication termination (Nicholls *et al*., 2018), TOP3A also removes the torsional stress during mtDNA replication (Hangas *et al*., 2022; Erdinc *et al*., 2023). It is perfectly plausible that due to the small size of the mitochondrial genome the nicking action for this function also occurs on the NCR, occasionally resulting in the breakage of the newly replicated part (Figure 4D), especially when H-strand replication is paused for a prolonged time at OriL (Torregrosa-Munumer *et al*., 2019). Although there are similarities with Mirin treatment and TOP3A knockdown, including the reduction of TOP3A protein levels (Figure 3B, Figure S4C), the knockdown caused accumulation of the OriH-OriL fragments in MGME1 KO cells and did not affect the 7S levels (Figure S4A), whereas Mirin had an opposite effect (Figure 1C, Figure 4A). As Mirin also does not appear to inhibit the ability of TOP3A to unwind supercoiled ssDNA (Figure S5), it is likely that TOP3A requires an additional interaction partner for the site-specific cleavage of mtDNA (Figure 4C–D). As of how Mirin can also influence the mtDNA supercoiling and the 7S DNA levels (Figure 4A) remains to be elucidated. However, it is likely that these effects are interconnected as one can imagine that reduced D-loop formation should increase mtDNA supercoiling. It can also well be that the relaxation activity of TOP3A is not the direct target for Mirin, as different pathological mutations in the enzyme can produce relatively different outcomes in terms of their molecular effects as well as pathology (Erdinc *et al*., 2023). Future research will hopefully decipher the inconsistencies in the reported Mirin targets as well as increase our understanding about the relationships of mtDNA replication termination, 7S DNA and mtDNA topology.

## Materials and Methods

### Cell culture

HEK293T, HEK293T *MGME1* knockout (Torregrosa-Munumer *et al*., 2019) and stable, inducible MTS-TOP3A-NLSmut-myc Flp-In^TM^ T-Rex^TM^-293 (Hangas *et al*., 2022) – hereafter mtTOP3A – cells were cultured in Dulbecco’s Modified Eagle Medium containing 4.5 g/l glucose, 2 mM L-glutamine, 1 mM sodium pyruvate, and 10 % fetal bovine serum at 37 °C in a humidified atmosphere with 7.5 % CO_2_. Expression of the transgene was induced by the addition of 5 ng/ml doxycycline to the growth medium for 48 hours. For mirin treatment, cells were seeded the day before the experiment, and subsequently 100 µM Mirin (Sigma-Aldrich) in DMSO was added directly into the growth medium for 48 hours. SiRNA transfections were carried out with Lipofectamine^TM^ RNAiMAX (Thermo Scientific) following the manufacturer’s recommended procedure, using 20 nM of siRNA and 21 µl of reagent, both in 1.5 ml of serum-free medium, per 10 ml plate. Dharmacon siGENOME SMARTpool human TOP3A siRNA (M-005279-01-0050) and siGENOME Non-Targeting siRNA Pool #1 (D-001206-13-05) were used for *TOP3A* knockdown and control transfections, respectively.

### Western blotting

To prepare whole-cell extracts for immunoblotting, cells were washed with 1 x PBS and resuspended in RIPA lysis buffer (25 mM Tris-HCl pH 7.6, 150 mM NaCl, 1 % NP-40, 1 % sodium deoxycholate, 0.1 % sodium dodecyl sulfate supplemented with protease and phosphatase inhibitor cocktails), the lysate was incubated on ice for 15 minutes followed by centrifugation at 14,000 *g* for 10 minutes at 4 ℃. Supernatant containing cellular proteins was transferred to a new tube and protein concentration was determined by Bradford assay, where bovine serum albumin (BSA) solution (Thermo Scientific^™^) was used for protein concentration reference standard. For immunoblotting, 50 µg of total protein were separated over an 8-12 % Tris/glycine SDS-PAGE, electrophoretically transferred to nitrocellulose membrane, blocked with 5 % non-fat milk in 1x TBST, incubated with antibodies, and detected using enhanced chemiluminescence (Bio-Rad ChemiDoc imaging system). Primary antibodies used were: rabbit anti-phospho-Stat1 (1:1000; #9167; Cell Signaling), mouse anti-β-tubulin (1:10 000; 66240-1-Ig; Proteintech), rabbit anti-TOP3A (1:1000; ABIN3181711; Antibodies-online), mouse anti-TWNK (1:1000; kind gift from Dr A. Suomalainen Wartiovaara), rabbit anti-MTCO2 (1:3000; 55070-1-AP; Proteintech), mouse anti-MRE11 (1:1000; ab214; Abcam), mouse anti-TFAM (1:10000; ab119684; Abcam), rabbit anti-TOMM40 (1:1000; sc-11414; Santa Cruz Biotech.), mouse anti-GAPDH (1:10000; 60004-1-Ig; Proteintech), mouse anti-Polδ (1:1000; 610972; BD Transduction Laboratories^™^). Secondary antibodies for immunoblotting were HRP-conjugated goat-anti-rabbit IgG (1:10000; A16104; Life Technologies) and goat-anti-mouse IgG (1:10000; ABIN101744; Antibodies-online).

### Gel electrophoresis

MtDNA topology and replication intermediates were analyzed by one- and two-dimensional agarose gel electrophoresis and Southern blotting (Goffart and Pohjoismaki, 2023). Briefly, cells were harvested, washed once with 1 x PBS and pelleted by centrifugation at 400 *g* for 4 minutes. Cells were resuspended in 500 µl of SDS lysis buffer (0.4 % SDS, 10 mM EDTA, 150 mM NaCl, 10 mM Tris-HCl, pH 7.4) supplemented with 100 µg of proteinase K and lysed overnight at 37 ℃. Upon lysis lipids and cellular debris (e.g., proteins) were separated from the DNA by addition of one volume of phenol:chloroform. Following centrifugation at 15,000 *g* for 5 minutes at room temperature (RT) an aqueous fraction containing DNA was transferred to a new tube. To remove traces of phenol an equal volume of chloroform was added to aqueous fraction and centrifuged again at 15,000 *g* for 5 minutes at RT. Final supernatant was carefully transferred to a new tube and the DNA was precipitated with 2.5 volumes of 100 % ice-cold ethanol for overnight at -20 ℃. Precipitated DNA was pelleted by centrifugation at 16,000 *g* for 30 minutes at 4 ℃ and the pellet was further washed with 500 µl of 75 % ethanol to remove salts. The supernatant was discarded and the DNA pellet was air-dried to remove any traces of ethanol. DNA was dissolved overnight at 37 °C in 50 µl of 1 x FastDigest buffer (Thermo Scientific^™^) in the presence of FastDigest *Bgl*II (Thermo Scientific^™^), which cuts the nuclear DNA but not mitochondrial DNA. DNA concentrations were determined by NanoDrop^TM^1000 spectrophotometer.

For mtDNA topology, 2 µg of uncut total DNA was separated over a 0.4 % agarose gel in 1 x TBE at 30 V without DNA dye for ∼16 hours at RT. Gel was then washed twice for 15 minutes with depurination buffer (0.25 M HCl) and twice for 20 minutes with denaturation buffer (0.5 M NaOH and 1.5 M) to improve the efficiency of DNA transfer. DNA was transferred overnight on positively charged nylon membrane (Amersham Hybond-XL) via moisture gradient by placing a pre-soaked membrane (in milliQ water) on the gel followed by a layer of Whatman filter paper and a stack of paper towels on top with ∼1 kg of weight. The next day the membrane was briefly rinsed with 6 x SSC (neutralization) and then baked at 80 ℃ for 2 hours to bind (crosslink) the DNA on the membrane. To detect for DNA containing complementary sequences on membrane-bound DNA a α-^32^P-dCTP-labelled probe spanning the region of 37-611 nts on human mtDNA was synthesized using High Prime DNA Labeling Kit (Roche). After pre-hybridizing the membrane with Church’s buffer (250 mM NaP_i_ pH 7.2, 7 % SDS, 2 mM EDTA) at 65 ℃ in a roller-blot hybridization oven for 20 minutes the buffer was changed for a fresh one (∼10 ml of Church’s buffer). Radioactively labelled probe was then added directly into the fresh buffer and the membrane was hybridized overnight at 65 ℃. On the next day the membrane was washed three times for 20 minutes with 1 x SSC and 0.1 % SDS, the membrane was then quickly dried and the signals were captured on phosphor screen (KODAK BAS). For mtDNA species analyses, 2 µg of total DNA was digested with 1 µl of FastDigest *Hind*III (Thermo Scientific^™^) for ∼6 hours at 37 °C, upon which half of the samples were treated with 1U (0.1 µl) of T7 Endonuclease I (New England Biolabs), which cleaves branched DNA structures, for 15 minutes at 37 °C, extracted once with phenol:chloroform and separated on a 0.8 % agarose gel in 1x TAE with 0.5 µg/ml of ethidium bromide (EtBr) at 30 V for ∼16 hours at RT. Southern blotting was carried out as before and the blot was hybridized with α-^32^P-dCTP-labelled 7S DNA (nts 16177-40) probe.

For 2D-gels, 10 µg of total DNA was digested with 4 µl of FastDigest *Hinc*II (Thermo Scientific^™^) for ∼16 hours at 37 °C, samples were then extracted once with phenol:chloroform and separated over a 0.4 % agarose gel in 1 x TBE without EtBr at low voltage (20-30 V), until the fragments of interest had migrated ∼10 cm into the gel. Upon electrophoresis the gel was stained with 1 µg/ml of EtBr and imaged. The lanes were cut out and moved onto a larger gel tray for the second dimension, and a 0.95 % agarose gel containing 1 µg/ml EtBr was cast around the first-dimension agarose slices. Second-dimension electrophoresis was run at 110 V in 1 x TBE buffer with 0.5 µg/ml EtBr at 4 °C until the linear fragments were ∼1 cm before the end of the gel. Southern blotting was performed as described before and the membrane was hybridized with a α-^32^P-dCTP-labelled probe for human mtDNA nucleotides 37-611. Radioactive signals were captured and quantified using phosphor screens and a phosphor imager (FLA-300, Fuji).

### Microscopy

For immunofluorescence, HEK293 *MGME1* knockout cells were grown on coverslips in a 6-well plate. The cells were fixed with 3.3 % paraformaldehyde in cell culture medium for 25 min, washed 3x with PBS and permeabilized with 0.5 % Triton X-100 in PBS/10% fetal bovine serum (FBS) for 15 minutes. Coverslips were incubated with the following primary and secondary antibodies in PBS/10 % FBS for 1 hour to overnight: mouse anti-MRE11 (abcam, #ab214, 1:500), rabbit anti-TFAM (antibodies-online, #ABIN2777277, 1:1000), mouse anti-DNA (abcam, #ab27156, 1:500), rabbit anti-TOMM20 (Sigma Life Sciences #HPA011562, 1:1000), Alexa Fluor 488 anti-rabbit IgG (Proteintech #SA00013-2, 1:1000) and Alexa Fluor 568 anti-mouse IgG (Lifetech, #A11032, 1:1000). After immunostaining, coverslips were mounted with ProLong^™^ Gold Antifade Mountant with DAPI (Thermo Scientific) and image acquisition was carried out using a Zeiss LSM710/LSM700 confocal microscope with 63x oil immersion objective.

### *In vitro* topoisomerase assays

Recombinant TOP3A was expressed and purified as previously described (Erdinc *et al*., 2023). The relaxation assay mixture contained 200 ng of supercoiled pUC19, 25 mM Tris-HCl pH 7.4, 50 mM NaCl, 5 mM MgCl_2_, 1 mM DTT, 0.1 mg/ml BSA, 30 nM TOP3A and the indicated concentration Mirin (Sigma #M9948) diluted in dimethyl sulfoxide. Reactions were performed at 37°C for 45 min and terminated by addition of 0.4% SDS. 6 × loading buffer (Thermo Scientific) was added and the reaction mixture was separated on a 1% agarose gel in 1x TAE (40 mM Tris, 1 mM EDTA, 20 mM acetic acid) at 120 V for 2 h. After electrophoresis, DNA bands were stained with 3 × GelRed and visualized on a Chemidoc imager (Bio-Rad).

### Gel mobility shift assay

DNA intercalation was studied by gel mobility shift assay, incubating 500 ng of supercoiled plasmid DNA (pcDNA5 FRT/TO and pOG44) with 100 µM Mirin (Sigma-Aldrich), 3.4 µM doxorubicin (known to exhibit strong DNA intercalation) (Sigma-Aldrich) and 0.5 µM 3,11-difluoro-6,8,13-trimethylquino [4,3,2-kl]acridinium methylsulfate (RHPS4) (Sigma-Aldrich). The plasmid DNA was separated over a 0.8 % agarose gel 1x TBE buffer without ethidium bromide for 2 hours at 90 V. The gel was then stained with 0.5 µg/ml of ethidium bromide and imaged using a Bio-Rad Chemidoc imaging system.

## Data Availability

This study includes no data deposited in external repositories and all the data are contained within the manuscript and supporting information file. The authors are willing to share the cell lines, reagents, laboratory notes and advice upon reasonable request for non-commercial purposes.

## Abbreviations

2D-AGE: Two-dimensional agarose gel electrophoresis
BTR complex: BLM-TOP3A-RMI1-RMI2 complex
DAMP: Damage-associated molecular pattern
DSB: Double-strand break
HR: Homologous recombination
mtDNA: Mitochondrial DNA
KO: Knockout
NCR: Non-coding region (of mtDNA)
OB-fold: Oligonucleotide-binding fold
OriH: Origin of heavy-strand replication
OriL: Origin of light-strand replication

## Acknowledgements

This study was funded by the Academy of Finland grants 325015 and 338227 to JLOP, and 332458 to SG. We are grateful to Prof. Anu Wartiovaara (Suomalainen) for providing us with the TWNK-antibody. We thank Ms. Anita Kervinen for her technical assistance in the laboratory, and Dr. Maria Falkenberg for insightful discussions.

