## Supplementary material for "A small molecule inhibitor Mirin prevents TOP3A-dependent mtDNA breakage and segregation"

**Running title: Mitochondrial effects of Mirin**

**Keywords:** MGME1 / mitochondrial DNA / DNA topology / DNA replication / double-strand break / DAMP / inflammation / molecular biology

### Supplementery Information

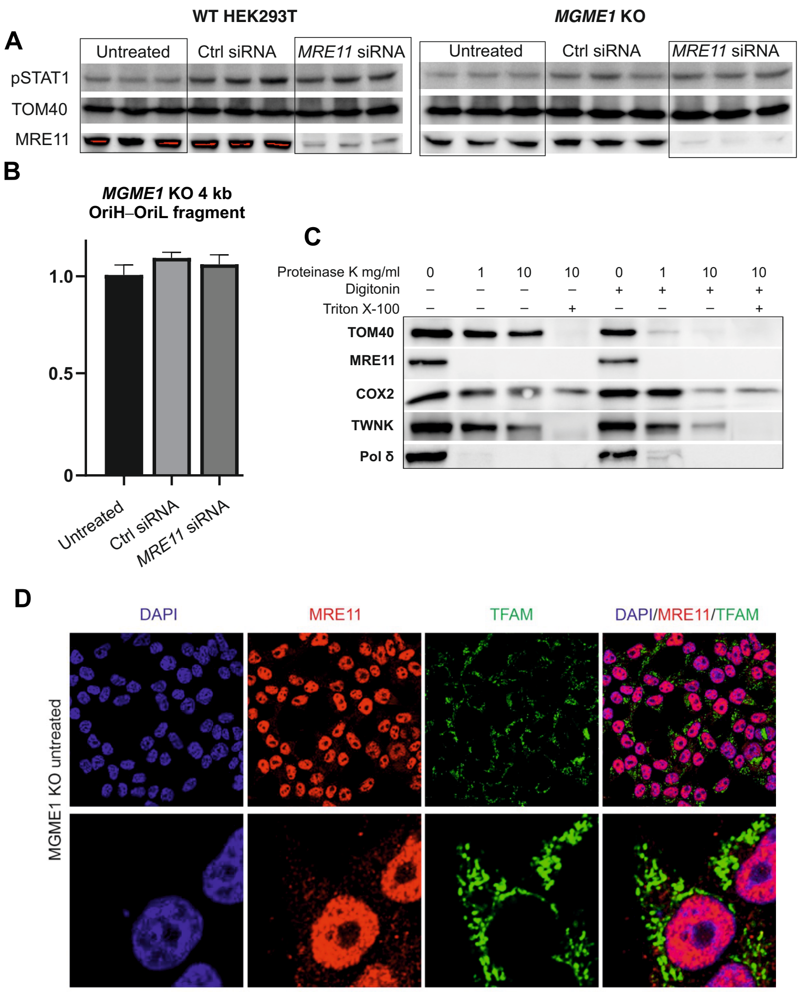


**Figure S1.** MRE11 is not localizing to mitochondria. (A) Knockdown of MRE11 in wildtype or MGME1 KO cells does not influence STAT1 activation or the abundance of OriH-OriL fragment (B). (C) In contrast to known mitochondrial proteins, such as TOM40, COX2 and TWNK helicase, MRE11 is not protected from protease treatment in mitochondrial preparations. Digitonin permeabilizes the outer membranes, rendering the outer membrane protein TOM40 sensitive to the proteinase treatment, while the matrix proteins COX2 and TWNK remain protected. Triton X-100 solubilizes mitochondria and exposes its proteins to the proteinase. Nuclear DNA polymerase δ is used as a nuclear protein contaminant control, being sensitive to proteinase already prior to detergent treatments. (D) Confocal fluorescent microscopy images of MRE11 localization in *MGME1* KO HEK293 cells. DAPI used for nuclear DNA and TFAM for mitochondrial localization. No colocalization of MRE11 with TFAM is evident. Bottom panels represent higher magnification of the upper.

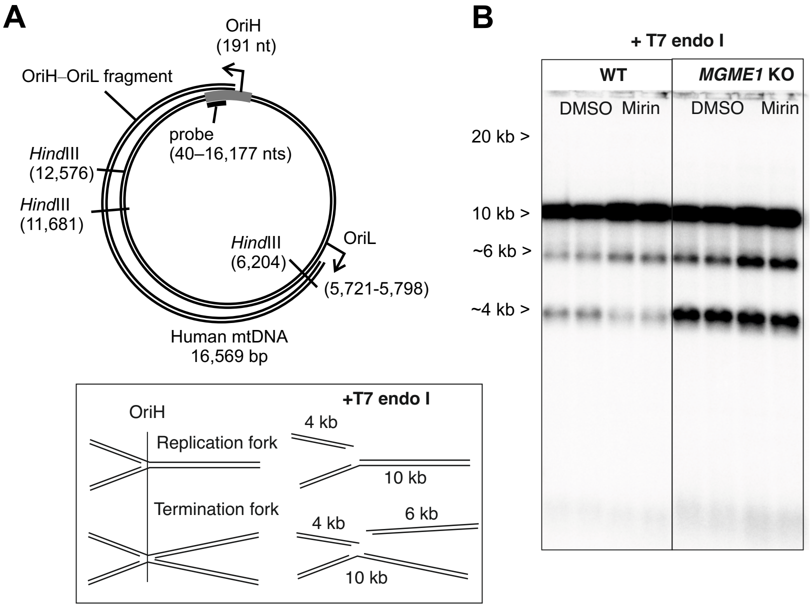


**Figure S2.** Effects of Mirin on mtDNA replication termination intermediates in the NCR. (A) Schematic illustration of mtDNA showing restriction cut sites, probe location and the extent of the OriH-OriL fragment of the MGME1 KO cells. Bottom panel: T7 endonuclease I (T7 endo I) cuts branched DNA and generates two distinct bands in *Hind*III-digested mtDNA, when probing for the 10 kb NCR containing fragment. The 4 kb fragment results from cleavage of both paused replication forks as well as from the termination intermediates, while the latter also produce a 6 kb fragment. (B) Mirin treatment changes the relative change in the intensities of 4 kb and 6 kb T7 endonuclease I digestion products. Note that while the 4 kb band is visible in the MGME1 KO cells without T7 endonuclease I digestion (Figure 1C), the 6 kb is not, indicating that the fragment is generated from some of the stalled intermediates evident in the 2D-AGE (Figure 2). Interestingly, the T7 endonuclease I-generated 6 kb fragment is detected by a probe situated downstream of OriH (compare with Figure 4C). Overall, the structure and exact location of the termination intermediates is poorly known and warrants further investigation.

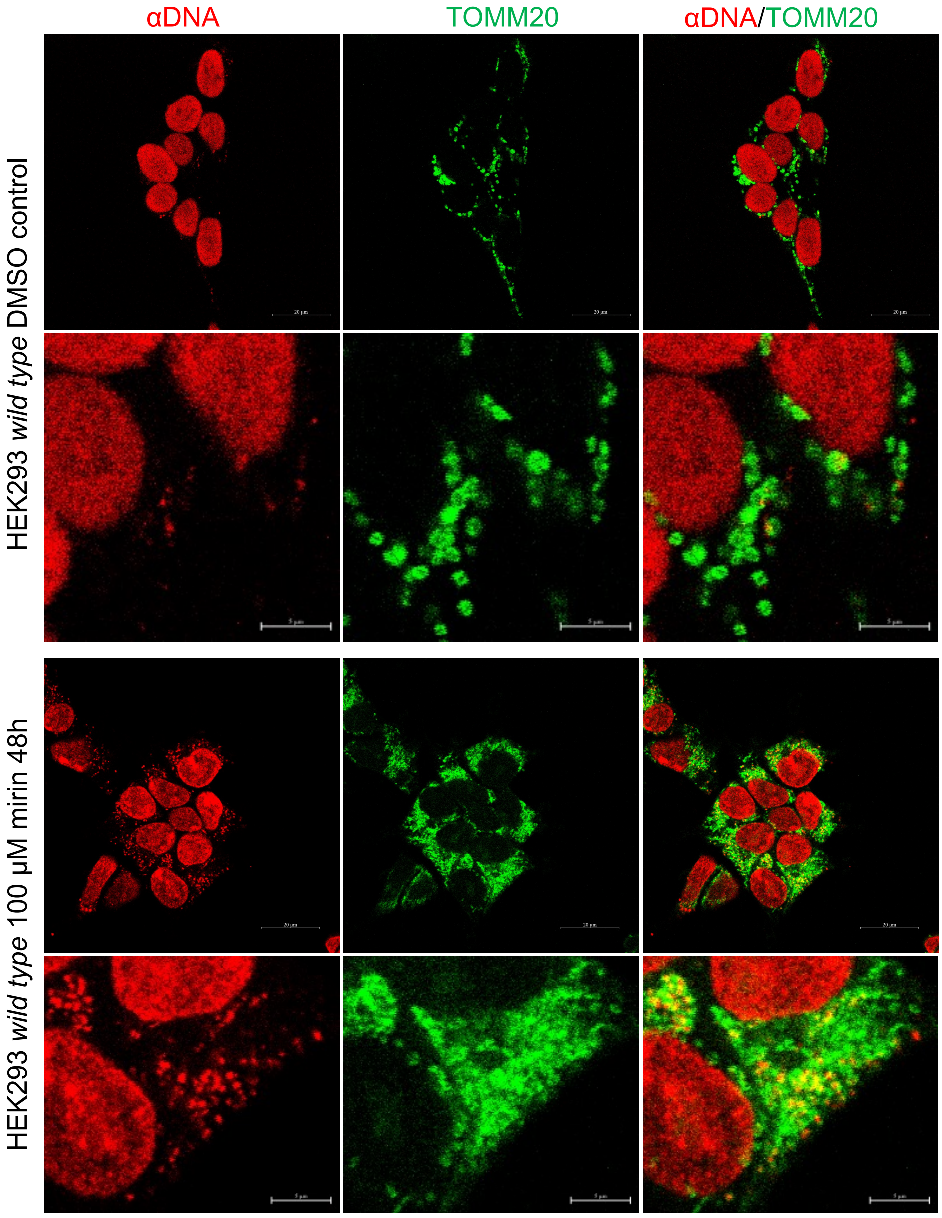


**Figure S3.** Mirin induces the aggregation of mitochondrial nucleoids. Top panels: Control HEK293 cells treated with DMSO for 48 h, stained with anti-DNA (red) and anti-TOMM20 (green). Lower panels: HEK293 cells treated with 100 μM Mirin for 48h.

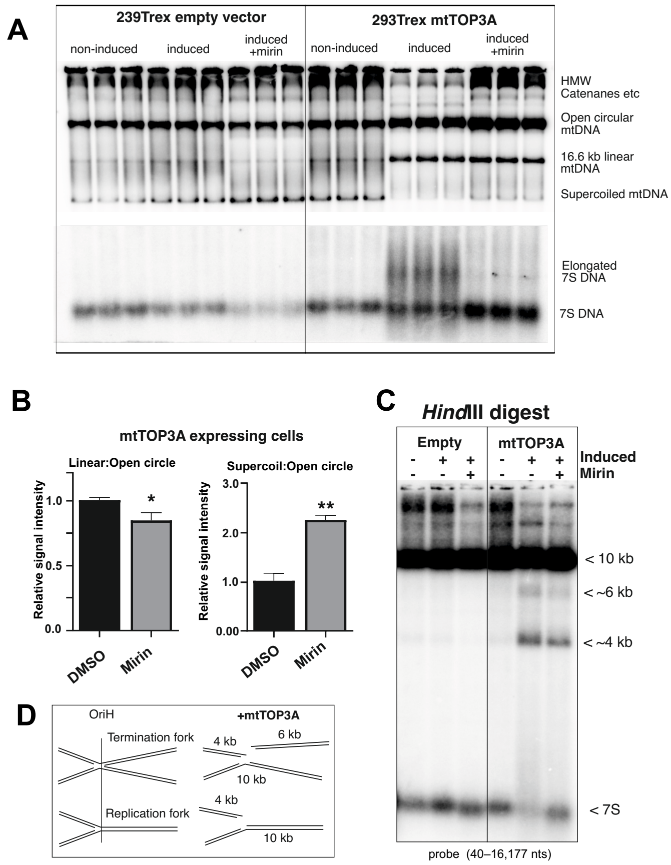


**Figure S4.** *TOP3A* knockdown increases the OriH-OriL fragment in *MGME1* KO cells. (A) Effects of TOP3A knockdown (kd) on the OriH-OriL fragment in wildtype (WT) and *MGME1* KO cells. Southern blot of HindIII-digested DNA probed for the 10 kb NCR containing fragment (see Figure 1B, C). Note the increase in 7S DNA in *TOP3A* kd WT cells, but not in *MGME1* KO cells, as well as the slight increase in the 4 kb OriH-OriL fragment in the latter. (B) Quantification of the 4 kb OriH-OriL fragment in the MGME1 KO cells after *TOP3A* kd. Ratios of 4 kb OriH-OriL fragment/10.2 kb fragment from three independent experiments is shown with standard error of the mean. Statistical significance (*P*-value 0.00194) was calculated with two-tailed Student's t-test. (C) Confirmation of the *TOP3A* knockdown and the antibody specificity on Western blot with HSP60 and EF-Tu as loading controls. Note the unspecific band below the *TOP3A* siRNA-sensitive species, which is also reduced by Mirin treatment (see Figure 3B).

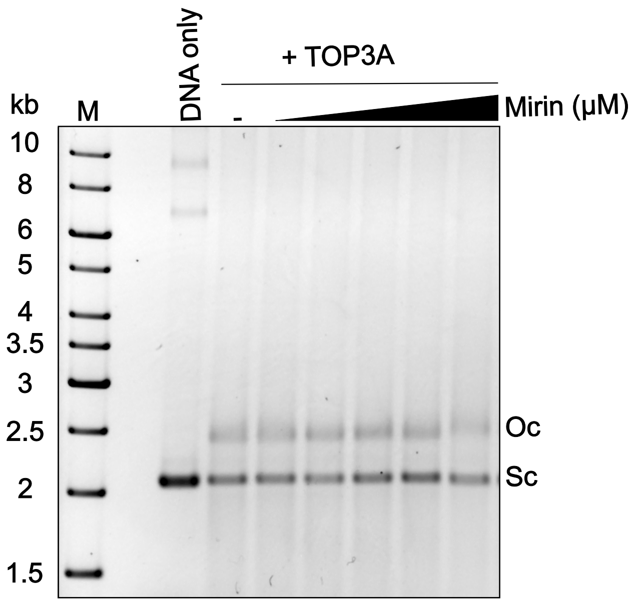


**Figure S5.** Mirin does not inhibit TOP3A supercoil relaxation activity *in vitro*. Purified TOP3A was incubated with 200 ng pBlueScript plasmid in the presence of increasing concentrations of Mirin (0.1, 1, 10, 100 and 1000 µM). The supercoiled plasmid (closed arrow) migrates faster in the electrophoresis than the open circles (open arrow).

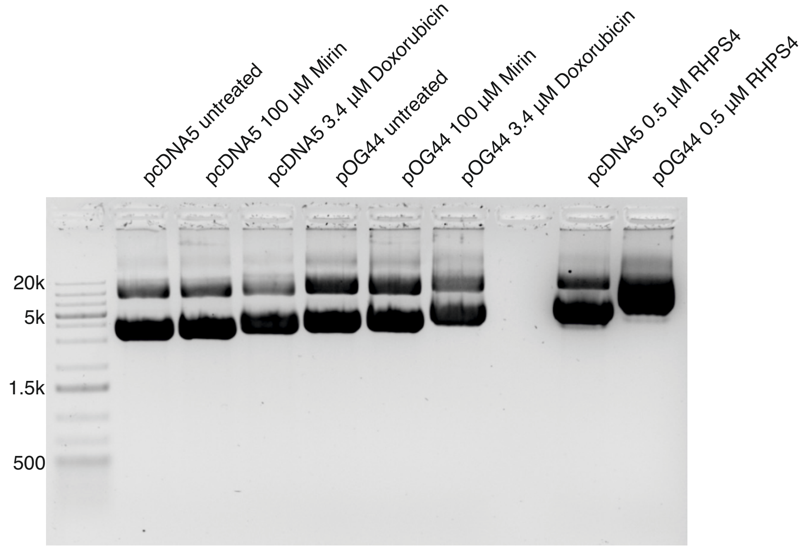


**Figure S6.** Mirin does not influence the electrophoretic properties of pcDNA5 and pOG44 plasmids *in vitro*. In contrast to the known DNA intercalators Doxorubicin and RSHP4, 100 µM Mirin had no effect on plasmid mobility in agarose gel electrophoresis.
